# EHE cell cultures: a platform for mechanistic and therapeutic investigation

**DOI:** 10.1101/2025.03.24.644191

**Authors:** Nicholas Scalora, Gillian DeWane, Yuliia Drebot, Ali A. Khan, Souradip Sinha, Krishnendu Ghosh, Denise Robinson, Patricia Cogswell, Andrew M. Bellizzi, Anthony N. Snow, Patrick Breheny, Michael S. Chimenti, Munir R. Tanas

## Abstract

Epithelioid hemangioendothelioma (EHE) is a difficult to treat vascular sarcoma defined by TAZ- CAMTA1 or YAP-TFE3 fusion proteins. Human cell lines needed to further understand the pathogenesis of EHE have been lacking. Herein, we describe a method to generate EHE extended primary cell cultures. An integrated multi –omic and functional approach was used to characterize these cultures. The cell cultures, relatively homogenous by single cell RNA-Seq, demonstrated established characteristics of EHE including increased proliferation, anchorage independent growth, as well as the overall gene expression profile and secondary genetic alterations seen in EHE. Whole genome sequencing (WGS) identified links to epigenetic modifying complexes, metabolic processes, and pointed to the importance of the extracellular matrix (ECM) in these tumors. Bulk RNA-Seq demonstrated upregulation of pathways including PI3K-Akt signaling, ECM/ECM receptor interaction, and the Hippo signaling pathway. Development of these extended primary cell cultures allowed for single-cell profiling which demonstrated different cell compartments within the cultures. Furthermore, the cultures served as a therapeutic platform to test the efficacy of TEAD inhibitors *in vitro*. Overall, the development of EHE primary cell cultures will aid in the mechanistic understanding of this sarcoma and serve as a model system to test new therapeutic approaches.

## Introduction

Epithelioid hemangioendothelioma (EHE) is a vascular sarcoma driven by specific chromosomal translocations that encode fusion genes involving *WWTR1* (TAZ is the protein) and *YAP1*. The predominant gene fusion is the *WWTR1*-*CAMTA1* fusion which joins the 5’ end of *WWTR1* (WW domain-containing transcription factor 1) in frame to the 3’ end of *CAMTA1* (calmodulin-binding transcription activator 1) to generate the TAZ-CAMTA1 fusion oncoprotein ^1,2^. This fusion is present in approximately 85-90% of EHE cases. The second fusion gene, *YAP1*-*TFE3*, involves the 5’ end of *YAP1* (Yes-associated protein 1) fused in frame to the 3’ end of *TFE3* (Transcription factor E3), creating a YAP-TFE3 fusion protein. This second fusion accounts for the remaining 10-15% of EHE cases ^3^. Recent reports of alternative fusion partners with *WWTR1* including *TFE3, MAML2*, and *ACTL6A* have also been identified in EHE ^4,5^. EHE cases can arise at almost any age, but typically occur between the ages of 30 and 40 with a slightly higher prevalence in females compared to males at a ratio of 1.1-1.5:1 ^6^. Tumors can appear in almost any organ in the body, but the most common sites for tumor formation are the liver and lungs ^7^. While surgery is the treatment of choice for unifocal EHE, there is no consensus on the best chemotherapies for progressing or metastatic EHE. Prior work has shown that over 50% of patients will develop metastatic EHE which impacts the overall survival of patients ^8^.

The Hippo signaling pathway is an evolutionarily conserved serine/threonine kinase cascade that regulates many cellular processes including cell growth, proliferation, apoptosis, and differentiation ^9–14^. The mammalian STE20-like protein kinases 1 and 2 (MST1/2) and large tumor suppressor 1 and 2 (LATS1/2) govern TAZ and YAP, the end effectors of the Hippo signaling pathway ^15–17^. TAZ and YAP are transcriptional co-activators that require binding the TEAD family of transcription factors in order to drive their transcriptional programs ^18,19^. Key upstream regulators of Hippo signaling include cell polarity, cell density, stress signals, and mechanical cues ^14,20,21^. Other known regulators of Hippo include LIFR, EGFR-MAPK, and LPS ^22–24^. Under sparse and attached conditions, TAZ and YAP are located within the nucleus where they can drive their transcriptional program. However, when cells become confluent or detached, TAZ and YAP are phosphorylated at key serine residues, shifting localization to the cytoplasm where they are ubiquitinated and degraded ^13,14,25^. The two fusion proteins that cause EHE dampen upstream negative regulation by LATS1/2, which is critical for their ability to remain within the nucleus and drive cellular transformation and anchorage-independent growth ^5,26–28^. Despite being composed of different C-terminal proteins, studies dissecting the mechanism of action of the TAZ-CAMTA1 and YAP-TFE3 fusion proteins revealed a similar mechanism of action. Bio-ID mass-spectrometry and subsequent functional studies showed that both fusion proteins interacted with the Ada-2a-containing histone acetyltransferase (ATAC) complex as well as other epigenetic regulators via the C termini of CAMTA1 and TFE3. These interactions facilitate TAZ-CAMTA1 and YAP-TFE3 driving similar transcriptional programs as assessed by principal component analysis of gene expression data ^27^.

In the absence of human EHE cell lines, the field has had to rely on engineered human cell lines and murine cell lines expressing the fusion proteins to gain insight into the pathogenesis of EHE. Both the NIH3T3 (spontaneously immortalized murine fibroblast cell line) and SW872 (human liposarcoma cell line), have been transduced to express either one of the two fusion proteins ^26–28^. New genetically engineered mouse models (GEMM) for the disease have been developed that express the TAZ-CAMTA1 fusion protein leading to formation of EHE ^29,30^. Crossing one of these GEMM models with a conditional *CDKN2A* knockout mouse, the most common secondary mutation in EHE ^31^, led to more aggressive EHE tumors in mice and the development of a mouse EHE cell line ^32^. More recently, a human PDX model of EHE has been developed ^33^. Despite these recent advances, the field still lacks human EHE cell line systems to complement the above approaches. Herein, we describe EHE extended primary cell culture systems that we anticipate can be used to validate previous findings while generating new hypotheses and therapeutic approaches.

## Results

### Generation of EHE extended primary cell cultures

To generate EHE cell cultures, EHE tumor samples were received from the EHE Biobank, an ethically-approved centralized biorepository after review and approval from central Independent Review Board. Of note, all samples were derived from female patients and originated in either the liver or the lung. EHE 036 is a metastatic sample, while the rest of the samples were derived from primary tumors **(Figure 1A)**. The clinical characteristics of the tumors utilized as well as the method of diagnostic confirmation are summarized (**Figure 1A**). The diagnosis of EHE was confirmed by histological analysis and a second method, either anchored multiplex PCR or immunohistochemistry for CD34 and CAMTA1 (**Figure 1A**). Histological analysis of EHE 036 and 053 show epithelioid (rounded) cells organized in cord-like structures and embedded in a myxohyaline matrix. Immunohistochemical expression of CAMTA1 supported the diagnosis of EHE **(Figure 1B)** as a surrogate for expression of the TAZ- CAMTA1 fusion protein, since *CAMTA1* is normally expressed predominantly in tissues of the central nervous system ^34^. The fresh or cryopreserved samples were obtained and grown in cell culture as shown (**Figure 1C**) and described in the **Materials and Methods** section, allowing cells migrating from the tumor to form monolayers in 2D culture (**Figure 1D**). In the image for EHE 036, a tumor fragment giving rise to tumor cells as shown is visible at the top of the image **(Figure 1D)**. While there were minor differences between each tumor, generally the adherent cells initially showed a rounded to plump spindle cell-shaped morphology. To validate the presence of the *WWTR1*-*CAMTA1* gene fusion in the extended primary cell culture, we performed RT-PCR followed by Sanger sequencing to identify the RNA breakpoint in EHE 001. EHE 001 shows exon 2 of *WWTR1* is fused in frame to the middle of exon 9 of *CAMTA1* creating an in-frame transcript **(Figure 1E)**. Anchored Multiplex PCR was performed on an EHE 036 tumor sample to identify the RNA breakpoint for EHE 036. The results showed that there are two isoforms of the fusion transcript separated by two codons including a more abundant transcript (**Figure 1F**) and a less frequent transcript (**Figure S1A, B**). Both of these fusion transcripts join the end of exon 2 of *WWTR1* in frame to a breakpoint in the middle of exon 9 of *CAMTA1* **(Figure 1E)**. We then designed a qRT-PCR assay characterized by a forward primer hybridizing to *WWTR1*, a reverse primer hybridizing to *CAMTA1*, and a fluorescence labeled custom probe spanning the breakpoint sequences for

**Figure 1.**
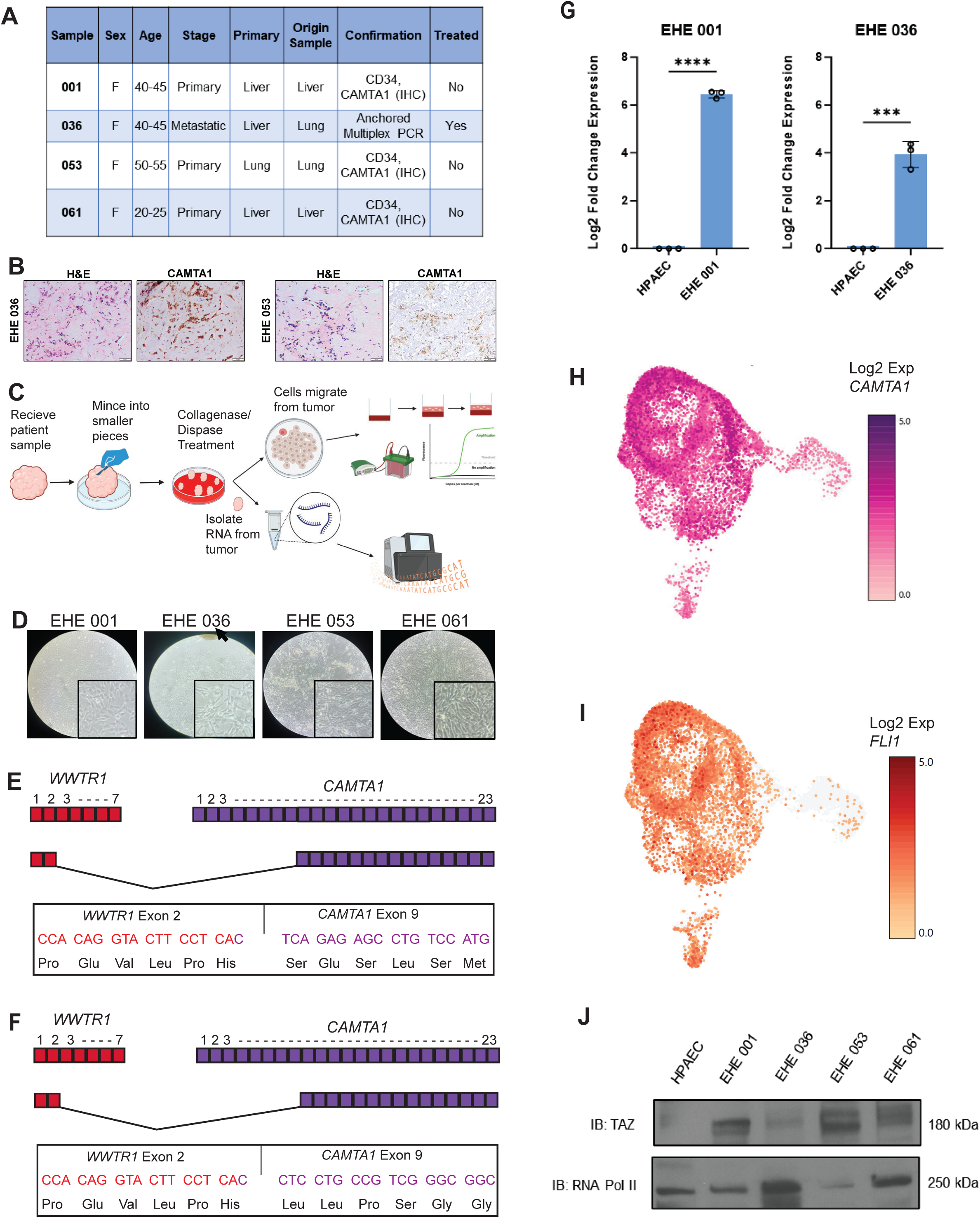
Derivation of EHE primary cell cultures and demonstration of the TAZ-CAMTA1 fusion protein (*WWTR1*-*CAMTA1* fusion gene). (**A**) Table containing patient characteristics associated with EHE tumors. (**B**) H&E staining and immunohistochemical evaluation of CAMTA1 expression in EHE 036 and EHE 053. Images taken at 200X. (**C**) Schematic outlining generation and characterization of the extended primary cell cultures. (**D**) Image of primary cell cultures in a monolayer with insets at higher power. Arrow denotes tumor fragment of EHE 036. (**E)** Schematic representation of RNA Breakpoint for EHE 001 determined by RT-PCR. Codons with their associated amino acids are shown below. (**F**) Schematic representation of RNA Breakpoint for EHE 036 by anchored multiplex PCR from tumor RNA. Codons with their associated amino acid are shown below. (**G**) qRT-PCR utilizing custom probes hybridizing to the EHE 001 and EHE 036 breakpoints respectively, with HPAEC as a control. (**H**) Single- cell RNA sequencing of early passage EHE 001 (passage number 6) demonstrating expression of *CAMTA1*. (**I**) Single-cell RNA sequencing of early passage EHE 001 (passage number 6) demonstrating expression of *FLI1*. (**J**) Western blot, after enriching for the nuclear fraction, probing for TAZ in HPAEC control and the EHE extended primary cell cultures. Error bars were used to define one standard deviation. For all panels, ****p<0.0001, *** p<0.001, ** p<0.01, *p<0.05.

EHE 001 and EHE 036. We see no expression of the fusion transcript in human pulmonary artery endothelial cells (HPAEC) control and do see amplification in the respective EHE culture **(Figure 1G)**. To determine the composition of these EHE extended primary cell cultures, we examined single cell RNA-Seq data of passage 6 (P6) cells using expression of *CAMTA1* as a surrogate for expression of the fusion gene. 76% of cells sequenced expressed *CAMTA1* **(Figure 1H)**; most of the cells negative for *CAMTA1* were in senescent cell populations demonstrating low transcriptional activity ^35^**, (Figure 5; Fig S5A-F)**. FLI1, a marker of endothelial differentiation was diffusely expressed in cells (**Figure 1I**) with the exception of senescent cells demonstrating reduced transcriptional activity (**Figure S5D**), consistent with the endothelial differentiation seen in EHE cells ^36^. To detect the TAZ-CAMTA1 protein, we enriched for nuclear protein by nuclear:cytoplasmic fractionation and probed with a TAZ antibody recognizing an epitope in the first two exons of TAZ, which are typically conserved in the fusion protein. Previous work has demonstrated TAZ-CAMTA1 fusion proteins with a molecular weight of approximately 175 kDa ^26^ that could vary slightly depending on the exact breakpoint. Western blot showed expression of a protein between 170-180 kDa in the four EHE extend primary cell cultures but not the human pulmonary aortic endothelial cells **(Figure 1J)**.

### EHE cell cultures show a transformed phenotype *in vitro* and are responsive to TEAD inhibitors

To determine if these extended primary cultures demonstrated a transformed phenotype, we performed various *in vitro* assays evaluating various hallmarks of cancer. All four EHE cell cultures showed increased proliferation compared to the HPAEC control by MTT-style proliferation assay **(Figure 2A)**.

**Figure 2.**
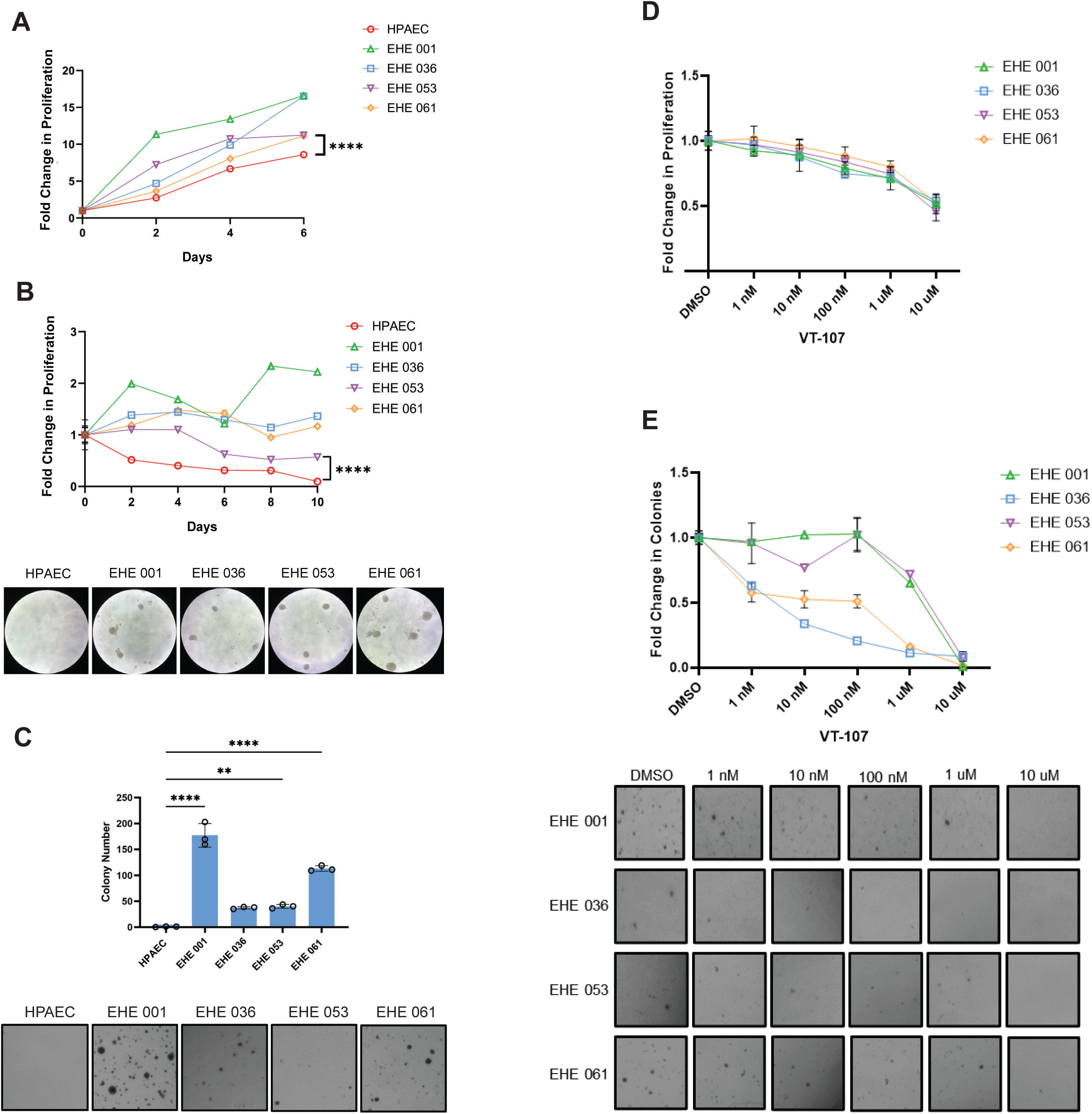
Functional characterization of EHE extended primary cell cultures. **(A**) Proliferation assay evaluating EHE extended primary cell cultures and human pulmonary artery endothelial cells (HPAEC). (**B**) Anchorage independent growth assay (poly-HEMA assay) with EHE extended primary cell cultures and HPAEC cells shown with corresponding images of spheroid formation below. (**C**) Soft agar assay with the same cells as in part A and B with corresponding images of soft agar colonies shown below (**D**) Proliferation dose response curve of the EHE extended primary cell cultures after treatment with pan- TEAD inhibitor VT-107. (**E**) Response of EHE extended primary cell cultures to treatment with VT-107 in soft agar with corresponding images below. For proliferation, poly-HEMA, soft agar assays, statistical significance was evaluated unpaired two-tailed Student’s *t*-test. Each experiment was repeated at least twice. Error bars were used to define one standard deviation. Smaller error bars do not extend beyond some individual data points on graphs. For all panels, ****p<0.0001, *** p<0.001, ** p<0.01, *p<0.05.

To test if these cells can grow in an anchorage independent manner, cells were grown on poly-HEMA (2- <u>h</u>ydroxy<u>e</u>thyl <u>m</u>eth<u>a</u>crylate) coated plates, a polymer that forces cells to grow in suspension. After 10 days, we observed no viable cells in the HPAEC control, in contrast with the EHE cultures which were still proliferating and forming spheroids. By day 10, EHE 001 was the most proliferative, followed by EHE 036 and EHE 061, then followed by EHE 053 **(Figure 2B)**. Anchorage independent growth was also assayed in soft agar. Consistent with the poly-HEMA assay, HPAEC cells did not proliferate in soft agar, while all EHE cell cultures formed colonies. Showing similar trends to the poly-HEMA assay, EHE 001 showed both the greatest number of colonies and the largest colonies of all EHE cultures, followed by EHE 061. EHE 036 and EHE 053 formed fewer and smaller colonies compared to EHE 001 **(Figure 2C)**.

The TAZ-CAMTA1 and YAP-TFE3 fusion proteins retain the TEAD binding domain of TAZ and YAP and rely on the TEA Domain (TEAD) family of transcription factors to drive their oncogenic transcriptome ^26,27^. TEAD inhibitors have recently been developed that disrupt the TAZ/YAP:TEAD interaction by inhibiting auto-palmitoylation of the TEAD transcription factors ^37,38^. To determine whether TEAD inhibitors would have efficacy in disrupting the TAZ-CAMTA1:TEAD interaction in EHE cells, and attenuate various hallmarks of cancer, EHE primary cell cultures were treated with VT- 107 (Vivace Therapeutics, San Mateo, CA), a pan TEAD inhibitor (inhibits TEAD1-4) ^39^. VT-107 inhibited proliferation in a concentration dependent manner across the four EHE extended primary cell cultures with a 30% decrease at 1 µM **(Figure 2D)**. To test the effect of VT-107 in EHE on anchorage independent growth, we performed a soft agar experiment with the drug at various concentrations. We observed that the primary cell cultures varied in their response to VT-107 with respect to anchorage independent growth **(Figure 2E)**. The EHE 001 and EHE 053 drug response curves largely mirrored each other as they were less sensitive to TEAD inhibition at lower concentrations but showed an approximately 20% decrease in colony formation at the physiologically achievable dose of 1µM, while EHE 036 and EHE 061 showed near 80% decrease in colony formation at this concentration. In contrast, IK-930 (Ikena Oncology, Boston, MA), a TEAD1-selective inhibitor ^40^ did not significantly affect proliferation (**Figure S2A**), but did inhibit anchorage independent growth in soft agar (**Figure S2B**).

### EHE extended primary cell cultures show low numbers of genomic alterations

The initiating genomic event in EHE are the chromosomal translocations and resultant gene fusions giving rise to the TAZ-CAMTA1 or YAP-TFE3 fusion proteins. However, additional genomic events have been implicated in the tumor progression of EHE. To determine whether these EHE extended primary cell cultures harbored similar genomic alterations, we performed whole genome sequencing on the four cell cultures (**Table S1, S2**). Initially, we examined whether our primary cell cultures demonstrated similar single nucleotide variants (SNV). From these set of genes, genes were identified that demonstrated one or two point mutations or splice alterations that were determined to impact disease progression in one of the EHE extended primary cell cultures ^7,31^ **(Figure 3A)**. We then examined the SNVs as a whole (**Table S1**) in an unbiased manner to gain further insight into the pathogenesis of EHE. Gene Ontology (GO) Cellular Component pathway enrichment analysis ^41^ showed enrichment for extracellular matrix-related ontologies (**Figure 3B**). Furthermore, genes within the mucin family demonstrated SNVs, a finding observed among all the primary cell cultures (**Figure 3C**). Comparison of copy number variants (CNV) demonstrated shared CNVs between many of the lines, as well as CNVs unique to each line **(Figure 3D, Table S2)**. GO Molecular Function pathway enrichment analysis ^41^ of the 34 shared CNVs showed enriched pathways related to metabolism via glutathione **(Figure 3E)**. Notable genes demonstrating CNVs have been highlighted (**Figure 3F**) due to their relevance to ECM interactions (*MUC3A*, *MUC12*, and *COL23A1*) an ontology also enriched in analysis of SNVs (**Figure 3B**) or glutathione metabolism (*GSTM1* and *GSTM2*) **(Figure 3E)** highlighting potential roles for these genes in the pathogenesis of EHE.

**Figure 3.**
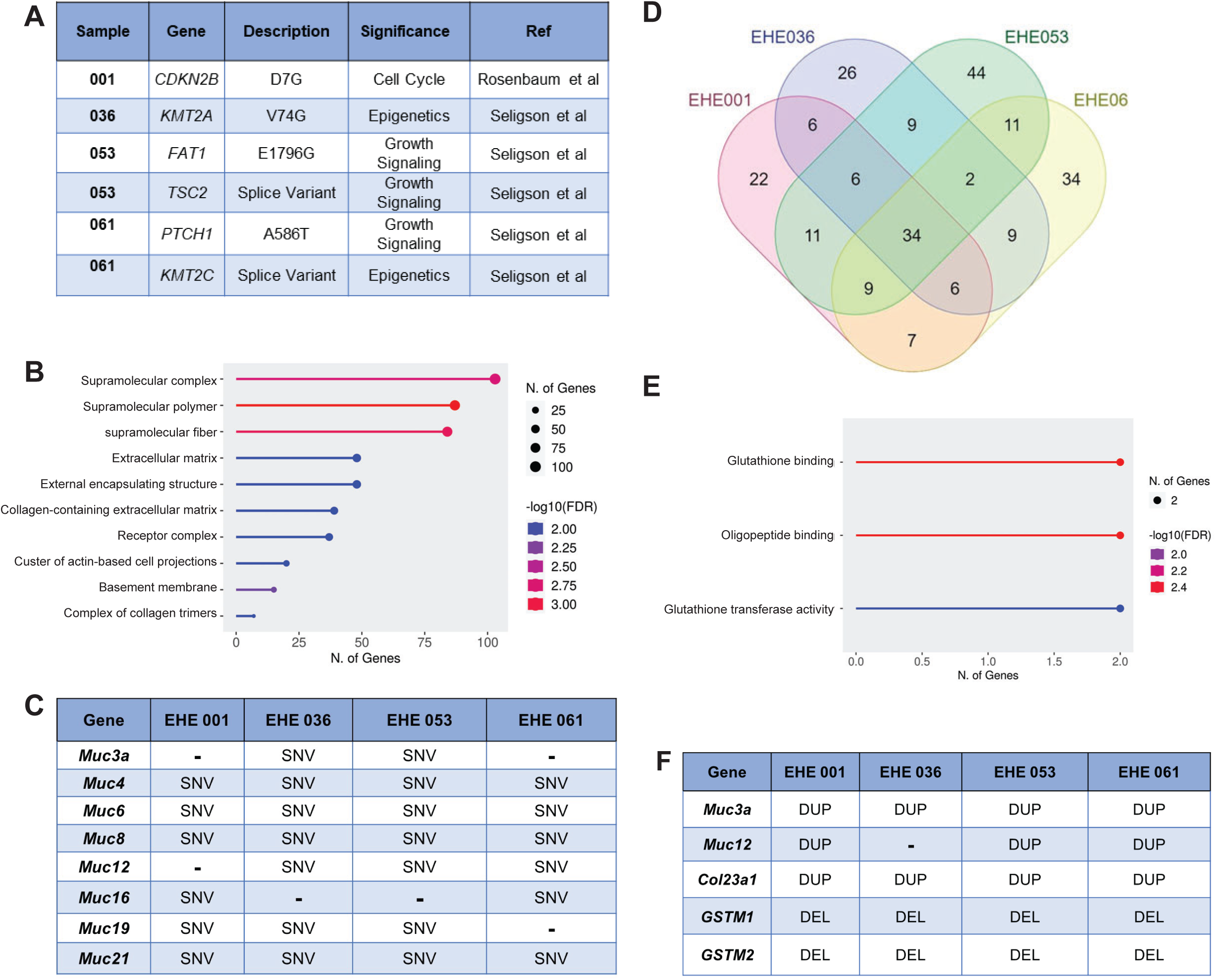
Genomic characterization of EHE extended primary cell cultures. **(A**) Table summarizing single nucleotide variants (SNVs) previously identified by whole genome sequencing in the EHE 001, EHE 036, EHE 053, and EHE 061 extended primary cell cultures and comparison with previous reports. Description of the amino acid substitution or other function consequence of the genomic alteration is included as well as the biological pathways involved. Similar genomic variants previously reported in the literature are represented. (**B**) GO Cellular Component pathway enrichment analysis of all moderate and high impact single nucleotide variants of the combined EHE cell cultures with an FDR of < 0.05. (**C**) Table describing the single nucleotide variants observed in the mucin genes in the EHE extended primary cell cultures. SNV indicates the presence of a single nucleotide variant. (**D**) Venn Diagram showing the overlap in copy number variants between all the extended primary cell cultures. (**E**) GO Molecular Function pathway enrichment analysis of the 34 shared copy number variants (CNV) among all EHE cell cultures with an FDR of < 0.05. (**F**) Table containing key genes in the CNV pathway analysis in panel (**E**). DUP indicates the presence of a duplication. DEL indicates the presence of a deletion.

### EHE primary cell cultures have similar transcriptomes

To further understand the transcriptomic profile of the extended primary cell cultures, we performed single cell RNA-Seq and bulk total RNA-Seq. According to our single-cell RNA sequencing saturation analysis, at a read depth of 20,000 reads per barcode (cell) we detected as many as 4,000 separate genes, with no significant plateauing of the line, consistent with a complex and diverse cancer transcriptome ^42^ **(Figure 4A)**. This is consistent with the hypertranscription phenotype caused by TAZ-CAMTA1 described previously in EHE ^43^. To further examine the transcriptomic profile among all the primary cell cultures, we performed bulk total RNA Sequencing including HPAEC controls and the donor tumor when available. By principal component analysis (PCA) the technical triplicates of EHE 001, 036, 053, and 061 clustered together, suggesting similar transcriptomes. At the same time, the primary EHE cultures demonstrated a transcriptome that differed from that of the HPAEC primary cells, indicating that the EHE cultures were expressing a transcriptome that did not simply recapitulate an endothelial cell signature (**Figure 4B**). Comparison of the EHE primary cell culture transcriptomes from the initial EHE tumors showed that the primary cell cultures drifted somewhat from the tumors they were derived from (**Figure 4B, C**). The transcriptomes from the extended primary cell cultures and EHE tumors (**Figure S3A, B**; **Figure S4A-D; Table S3**) were combined and compared (**Figure 4D, E; Figure S4E, F**). Overall, upregulated genes in the EHE cell cultures showed a greater degree of overlap (Matthew’s correlation coefficient [MCC] = 0.543; p < 2.2 x 10^-16^) (**Figure S4E**) than downregulated genes (MCC= 0.082; p < 2.2 x 10^-16^) (**Figure S4F**). Despite these differences, iPathway guide analysis (Advaita Bioinformatics, Ann Arbor, MI) of the combined transcriptomic profile of the donor tumors (**Figure 4D**) and EHE cell cultures (**Figure 4E**) showed that key cancer pathways including PI3K-Akt signaling, ECM receptor interaction, Hippo signaling pathway, proteoglycans in cancer, and focal adhesion were conserved, indicating that the EHE extended primary cell cultures recapitulate key components of the transcriptomes seen in EHE tumors.

**Figure 4.**
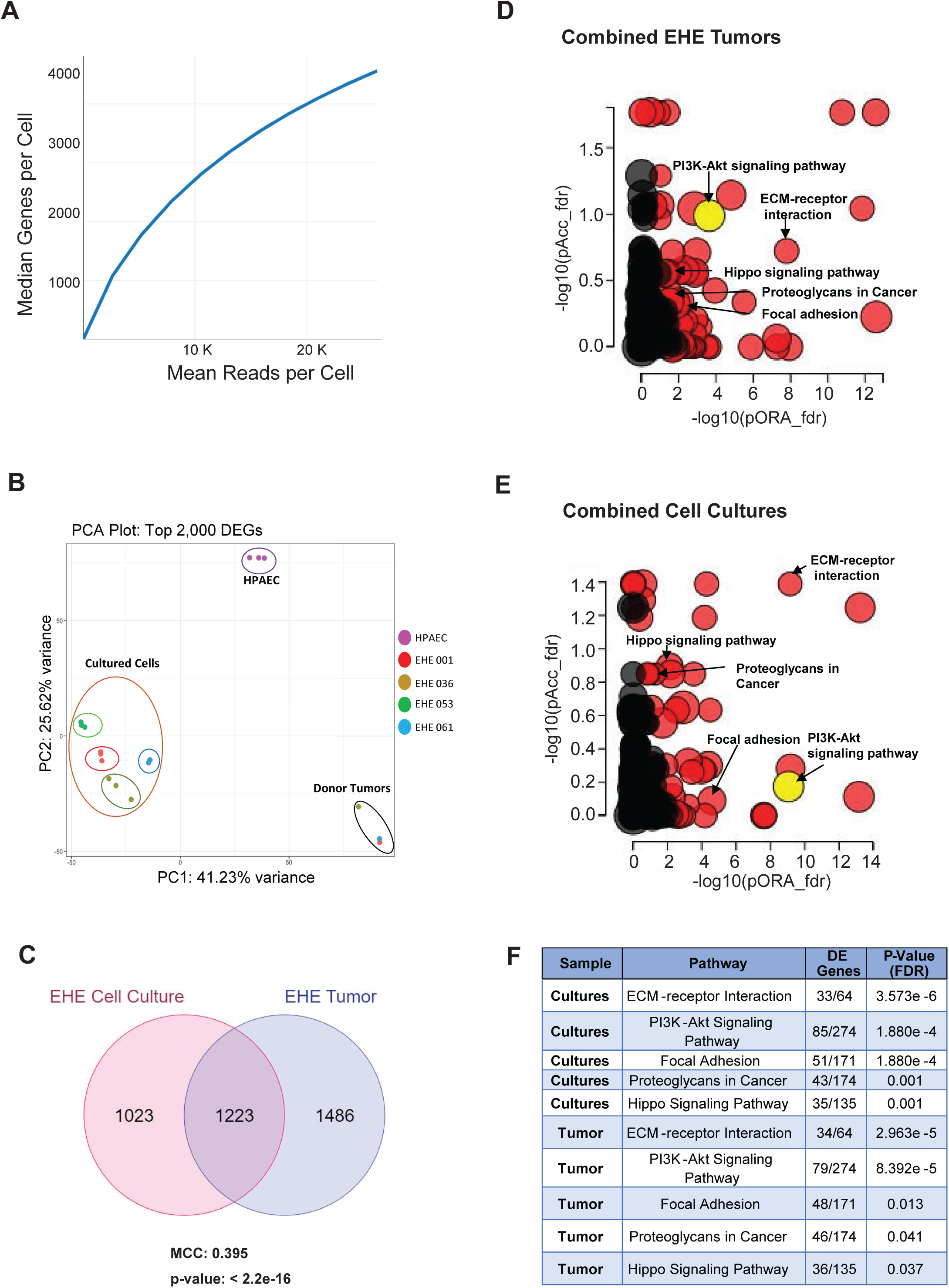
Transcriptomic profiling of EHE extended primary cell cultures. **(A**) Sequencing saturation of single cell RNA-seq for passage number 6 of EHE 001 (Median Genes per Cell vs. Mean Reads per Cell). (**B**) Principal component analysis of total RNA sequencing from the EHE tumors, the derivative extended primary cell cultures, and HPAEC control cells based on the top 2,000 differentially expressed genes (DEGs). (**C**) Venn diagram showing the overlap in differentially expressed genes between the EHE tumors and the combined EHE primary cell cultures. To assess enrichment, hypergeometric analyses and Matthews correlation coefficient (MCC) calculations were carried out in R, version 4.4.1. The upper tail of the hypergeometric distribution was calculated using the phyper() function. (**D**) Pathway enrichment analysis (iPathwayGuide analysis, Advaita Bioinformatics) for EHE tumors (transcriptomes are combined) plotting -log10 probability of accumulation (pAcc) vs probability of over- representation (pORA). (**E**) Pathway enrichment analysis (iPathwayGuide analysis, Advaita Bioinformatics) for EHE extended primary cell cultures (transcriptomes are combined). (**F**) Table further detailing the fraction of differentially expressed (DE) genes in enriched pathways in the EHE extended primary cell cultures and tumors.

**Figure 5.**
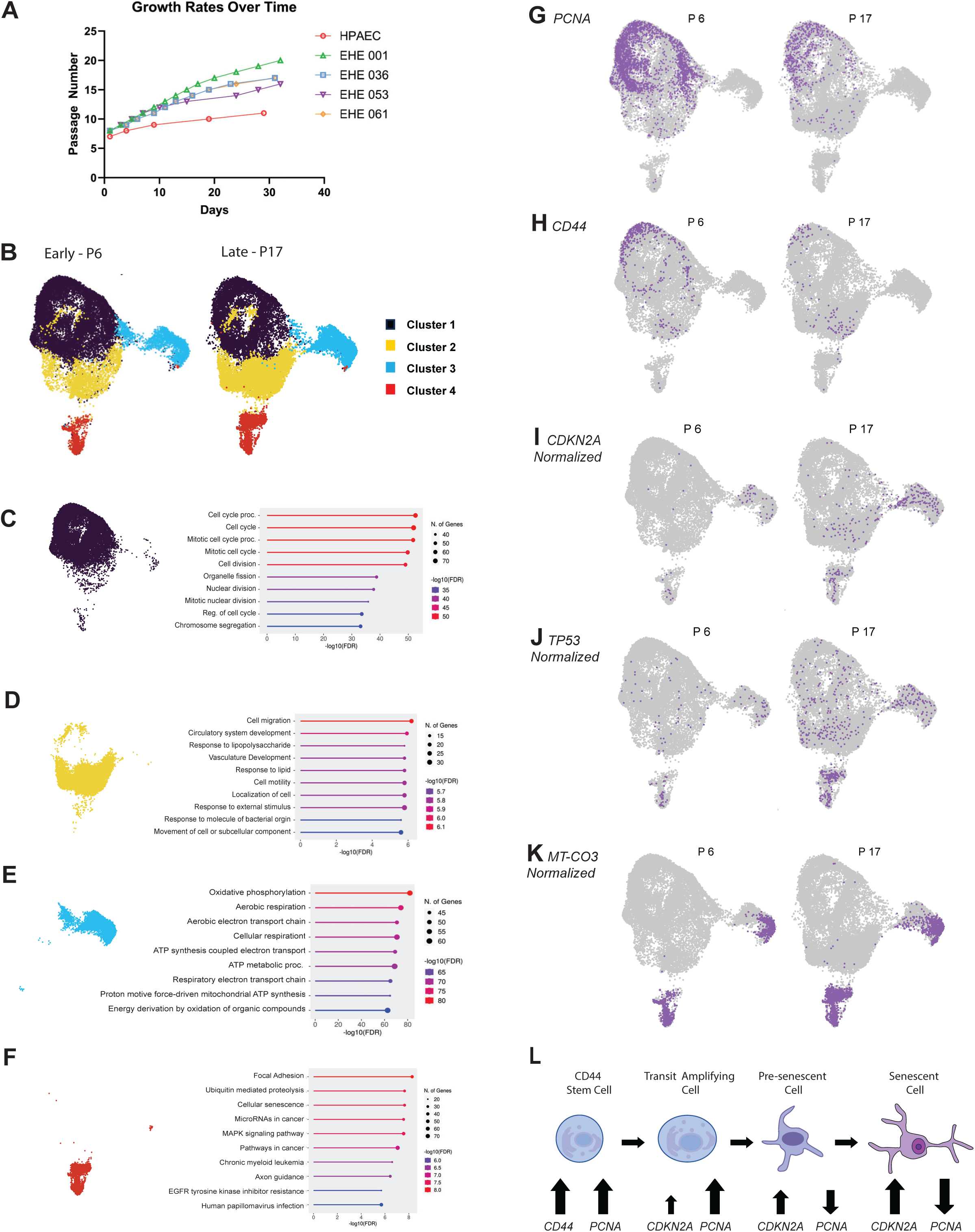
Composition of early and late passage EHE extended primary cell cultures. (**A**) Plot showing the growth rate of the EHE extended primary cell cultures as a function of days passaged. Data points on the line graphs represent passages. Human pulmonary artery endothelial cells (HPAECs) are included as a control. (**B**) t-distributed stochastic neighbor embedded (t-SNE) plot showing unbiased clustering of EHE 001 cells by Loupe browser. (**C**) GO Biological Process pathway enrichment analysis for Cluster 1. (**D**) GO Biological Process pathway enrichment analysis for Cluster 2. (**E**) GO Biological Process pathway enrichment analysis for Cluster 3. (**F**) Kyoto encyclopedia of genes and genomes (KEGG) analysis for Cluster 4. Identical thresholding for relative gene expression maintained for early and late passaging and was also maintained for the comparisons in (**G-K**). (**G**) Comparison of number of *PCNA* positive cells (purple) between P6 and P17. (**H**) Comparison of *CD44* positive cells between P6 and P17. (**I**) Comparison of *CDKN2A* positive cells normalized by transcriptional activity (UMI/cell count) between P6 and P17. (**J**) Comparison of *TP53* positive cells normalized by transcriptional activity (UMI/cell count) between P6 and P17. (**K**) Comparison of *MT-CO3* positive cells normalized by transcriptional activity (UMI/cell count) between P6 and P17. (**L)** Working model for different cell subsets in EHE extended primary cell cultures.

### Composition of the EHE primary cell cultures

As cells were continually passaged in an effort to obtain an immortalized cell line, an increase in the number of days between each passage and concurrent decrease in the proliferation of the extended primary cell cultures was noted, demonstrating the Hayflick principle associated with senescent cells ^44,45^ (**Figure 5A**). We observed little changes in morphology between P6-P17, but did observe increased cell size, binucleated cells, and cytosolic process (**Figure S5A**), increased expression of β-galactosidase by P20 consistent with senescence **(Figure S5B)**, and increased expression of p16 with continued passaging of the EHE primary cell cultures (**Figure S5C**). By the single cell RNA-Seq data, we found the there was a reduction in unique molecular identifiers (UMI)/cell in later passage cells (**Figure S5D, E**), consistent with the transcriptional dysregulation seen in aging/senescent cells ^46^. We then returned to examining the single cell RNA sequencing performed on the EHE 001 extended primary cell culture to determine if there were changes in various subsets of the extended primary cell culture that could explain the senescent changes.

To further evaluate the composition of the primary cell cultures, we evaluated unbiased clustering performed by the Loupe browser which showed that the EHE 001 extended primary cell culture clustered into four distinct groups by t-distributed stochastic neighbor embedded (t-SNE) plots (**Figure 5B**); cluster 1 (black), cluster 2 (yellow), cluster 3 (blue), and cluster 4 (red). To characterize the genes driving the clustering of cluster 1, we evaluated genes showing a 2-fold change and p < 0.05 (**Figure 5C**). 124 such genes were identified (**Table S4**). We then performed GO Biological Process pathway enrichment analysis ^41^, these genes were enriched for those involved in cell cycle progression and cell division (**Figure 5C**). Using similar thresholds for cluster 2 (**Figure 5D**), 77 such genes were identified (**Table S4**). Characterization of these genes by GO Biological Process pathway enrichment analysis showed they were enriched for cell migration, a hallmark of cancer. In addition, enrichment for vasculature development was also noted, consistent with cells in this cluster demonstrating endothelial cell differentiation (**Figure 5D)**. The threshold for genes evaluated in cluster 3 (**Figure 5E**) was log_2_(0.5) or 1.4-fold due to a fewer number of genes being differentially expressed. At this threshold, 134 genes were identified. GO Biological Process pathway enrichment analysis (**Figure 5E**) shows an enrichment for processes consistent with oxidative phosphorylation consistent with upregulation/dysregulation of mitochondrial genes (**Table S4**), often seen in senescent cells ^35,47^. The threshold for genes evaluated in cluster 4 (**Figure 5F**) is similar to that in clusters 1 and 2; 1480 genes were identified. Kyoto encyclopedia of genes and genomes (KEGG) analysis ^41^ showed several pathways enriched including genes associated with cellular senescence (**Figure 5F; Table S4**).

In order to determine potential mechanisms driving senescence, gene expression in the t-SNE plots of P6 and P17 cells were compared. *PCNA*, a marker of proliferation ^48^ was enriched in cluster 1, and generally absent in clusters 3 and 4, suggesting the clusters 3 and 4 were composed of a senescent population of cells. *PCNA* expression overall, however, was decreased in P17 cells vs. P6 cells (**Figure 5G**). Similar trends were observed for MKI-67, another known marker of proliferation ^49^ **(Figure S5F, G)**. Analysis of combined bulk RNA-Seq of the EHE extended primary cell cultures showed that *CD44* and *LIF*, cancer stem cell markers ^50,51^, were upregulated (**Figure S3A**) suggesting that a stem cell compartment might be present with these extended EHE primary cell cultures. To determine if this proliferative compartment coincided with the presence of a regenerative, stem-cell like compartment, we evaluated expression of *CD44* and *LIF*, and showed a cluster of *CD44* expressing cells (**Figure 5H**) concurrently showing higher expression of levels of *PCNA* (**Figure 5G)** and *MKI-67* **(Figure S5F, G**) in cluster 1. This *CD44* positive population is eventually exhausted by the P17 passage (**Figure 5H**). As similar pattern was noted for expression of *LIF* (**Figure S5H, I**). Together these findings led us to hypothesize that a regenerative cancer stem-cell like compartment exists in these EHE extended primary cell cultures that replenishes the actively proliferating compartment, and that eventually undergoes senescence with enough passages.

To explore this possibility further, we evaluated expression of two genes classically know to be upregulated in cellular senescence *CDKN2A* and *TP53* ^47^ . *CDKN2A* (**Figure 5I**) and *TP53* (**Figure 5J**) expression levels were both elevated in P17 cells as compared to P6 cells, consistent with previous findings indicating a trend towards senescence with increased passage number. Furthermore, when normalizing expression of the individual genes to the level of overall transcriptional activity, we noticed that *CDKN2A* and *TP53* levels were elevated in the two hypoproliferative clusters presumed to represent senescent cells, clusters 3 and 4 (**Figure 5E, F**) further supporting the hypothesis that these two smaller clusters represent senescent cells. Expression of mitochondrial genes such as *MT-CO3* described as markers in senescence and aging ^35,47,52^ were also elevated in clusters 3 and 4 **(Figure 5K)**. All together, the above findings suggest a working model (**Figure 5L**) consisting of a proliferative, stem cell-like compartment (cluster 1), that matures into a transit amplifying-like cell with a transcriptomic profile demonstrating endothelial differentiation and hallmarks of cancers such as migration (cluster 2).

Eventually *CDKN2A* and *TP53* expression is increased along with various mitochondrial genes consistent with pre-senescent and senescent cells (clusters 3 and 4).

## Discussion

### EHE cell cultures demonstrate key features of human EHE tumors

Despite the recent development of several *in vitro* and *in vivo* models of EHE, human EHE cell line systems have been limited to engineered cell lines systems. Here, we have identified a method for the generation of EHE extended primary cell cultures from human tumor samples that can be used to advance understanding of disease progression. The EHE extended primary cell cultures recapitulate key features of epithelioid hemangioendothelioma. The cell cultures demonstrate the TAZ-CAMTA1 fusion protein (*WWTR1*-*CAMTA1* gene fusion) by western blot, RT-PCR, and qRT-PCR. Anchored multiplex PCR for sample EHE 036 demonstrated two possible splicing isoforms. The determined RNA breakpoints are consistent with previously described *WWTR1*-*CAMTA1* fusion genes that demonstrate the end of exon 2 or 3 of *WWTR1* fused in frame with exon 8 or 9 of *CAMTA1* ^1,2,53^. Generating cultures from primary tumors often raises concerns regarding the heterogeneity of the cell culture. Our extended primary cell cultures show limited heterogeneity by passage number 6 with most cells lacking expression of *CAMTA1* limited to the non-proliferative, senescent population (**Figure 5E, F**), that shows reduced transcription overall (**Figure S5D, E**). Future generation of additional EHE extended primary cell cultures is warranted for the following reasons. First, there is evidence that extended primary cell cultures that are a few passages out from the primary tumor can recapitulate disease progression more accurately than immortalized cell lines that have been passaged for a much longer period of time, so generating multiple cell cultures for a cancer can be beneficial for mechanistic work and drug discovery ^54^. Furthermore, additional attempts to develop EHE extended primary cell cultures are needed in order to generate extended primary cell cultures from tumors expressing YAP-TFE3.

These extended primary cell cultures recapitulate other key features of epithelioid hemangioendothelioma. We have shown that EHE extended primary cell cultures proliferate faster compared to primary endothelial cell controls. These cultures also maintain the well-established phenotype of the TAZ-CAMTA1 fusion protein of anchorage independent growth demonstrated by growth on poly-HEMA and in soft agar ^26,27^. The cell cultures also exhibit reduced anchorage independent growth and to a lesser degree, proliferation, due to TEAD inhibition. The different cell cultures vary in their sensitivity to TEAD inhibition; future studies focused on understanding what might be driving the different therapeutic responses of the various EHE tumors/cell cultures is warranted. In addition, the EHE cell cultures differed in their responses to two different TEAD inhibitors, suggesting these extended primary cell cultures might represent platforms that can be used to determine the efficacy of various therapeutic approaches.

### EHE extended primary cell cultures show genomic alterations in epigenetic modifiers and ECM proteins

We have demonstrated that there are low numbers of genomic alterations in the primary cell cultures of EHE, in line with the established literature. For comparison, lung cancer has a high somatic mutation rate with an average of 8.1 mutations per mega base of DNA. On the lower end, acute myelogenous leukemia has on average 0.56 mutations per mega base of DNA^55^. In comparison, the EHE samples in this study displayed an average of 0.29 mutations per mega base. Notably, we saw SNV’s in *KMT2A* and *KMT2C* in different EHE cell cultures. The Compass-like complex, of which these genes are a part of, are known interactors of the TAZ-CAMTA1 and YAP-TFE3 fusion proteins ^27^. Additional studies are warranted to understand the functional significance of these SNVs and how recruitment of these epigenetic complexes plays a role in the progression of EHE.

Mutations in mucins identified in this study have not previously been characterized in EHE but might play an important role in disease progression since the tumor has a prominent extracellular matrix (ECM). Histologically, the neoplastic cells of EHE are embedded as single cells and cords in an abundant myxohyaline matrix. Previous studies using human cell line systems have shown that ECM-related genes are a key component of the transcriptional programs of TAZ-CAMTA1 and YAP-TFE3^27^ and were also identified by pathway analysis of bulk transcriptomic data in this current study (**Figure 4D-F** and **S4A- D**). Aberrant expression of mucins has been shown to play a role in masking cancer cells from immune surveillance and can aid in metastasis ^56,57^. Additionally, mutations in mucins portend a worse prognosis among many cancer types ^58^. All together, these observations warrant future studies elucidating the contributions of mucin and other extracellular matrix proteins in the pathogenesis of EHE. Enrichment of CNV alterations in glutathione metabolism pathways combined with single cell RNA-Seq data showing high expression of mitochondrial genes in a subset of cells which indicate a potential therapeutic vulnerability and also warrant future investigation.

### EHE tumors and primary cell cultures have similar transcriptomic profiles

As expected, the overall transcriptome of the EHE extended primary cell cultures differed somewhat from the tumors they were derived from. However, critical pathways that have been established as targets of the TAZ-CAMTA1 fusion protein in previous literature were conserved between tumor and the extended primary cell cultures. These include the PI3K-Akt signaling pathway, Hippo signaling pathway, and focal adhesion-related proteins ^27^. The extended primary cell cultures recapitulate other features of EHE including the hypertranscription phenotype. Recent work showed that exogenous expression of the TAZ- CAMTA1 fusion protein in endothelial cells led to hypertranscription and subsequent senescence, further highlighting similarities observed in our primary cell cultures and other findings in the field ^43^.

### EHE primary cell cultures show different compartments

In many ways, the prominent extracellular matrix of EHE mentioned above has made it challenging to isolate tumor cells and generate cell cultures. This in turn, has made EHE recalcitrant to evaluation of different cellular compartments within the tumor. Utilization of primary cell cultures devoid of extracellular matrix in this study has facilitated single cell RNA sequencing approaches. This approach has identified a *CD44, LIF* positive proliferative stem cell-like compartment that gives way to more mature transit amplifying-like cells demonstrating a transcriptomic profile reflecting vascular development and enrichment for genes involved in migration. Finally, senescent populations of cells can be seen characterized by increased *CDKN2A* and *TP53* expression.

### EHE extended primary cell cultures undergo senescence

The Hayflick principle describes the observation that normal cultured cells become senescent after a limited number of divisions, typically around 40-50 doublings ^44^. Although EHE is a malignant neoplasm, we did observe a senescent phenotype after serial passaging, consistent with replicative senescence and the shortening of telomeres. A study trying to derive a variety of sarcoma cell lines from patient samples showed that 40 of 47 cultures either never formed monolayers, or were senescent by passage number 10, indicating this phenomenon can be observed in other cancer types ^59^. Our data implicate loss of *TP53* and *CDKN2A* in this population, which is consistent with previous studies showing: frequent loss of *CDKN2A* in clinical samples ^7,31^, that *CDKN2A* loss results in more aggressive EHE *in vivo*, and that *CDKN2A* loss is also required to generate a murine cell line from the tumors ^32^. Future work generating additional lines increases the likelihood of a spontaneously immortalized cell line. Alternatively, deletion/inactivation of *CDKN2A* could lead to generation of immortalized EHE cell lines.

## Acknowledgements

This work was supported by a University of Iowa Sarcoma Multidisciplinary Oncology Group pilot award (M.R.T), by a predoctoral fellowship from the EHE Foundation (N.S), by a grant from the Veterans Health Administration Merit Review Program 1 I01 BX003644-01 (M.R.T), by grants from the National Institutes of Health, National Cancer Institute 1 R01 CA237031-01A1 (M.R.T), by an award from the University of Iowa Stead Family Scholars Program (M.R.T), and by an NCI Core Grant P30 CA086862 (University of Iowa Holden Comprehensive Cancer Center). Next generation sequencing data presented herein were generated by the Genomics Division of the Iowa Institute of Human Genetics (RRID: SCR_023422) which is supported, in part, by the University of Iowa Carver College of Medicine. We also want to acknowledge The EHE Foundation Biobank / Cleveland Clinic Central Biorepository for providing EHE tumor specimens for these studies.

## Author contributions

Conceptualization, M.R.T.; Methodology, A.M.B., P.B ,M.S.C.,., and M.R.T.; Software, M.S.C., P.B.; Formal analysis, A.N.S., M.S.C., P.B., and M.R.T.; Investigation, N.S., G.D, Y.D., A.K., S.S, K.G., A.M.B, A.N.S., P.B., M.S.C., and M.R.T. Resources, D.R., P.C., P.B., and M.R.T.; Writing-Original Draft, N.S.and M.R.T.; Writing-Review & Editing, N.S, G.D.,.and M.R.T.; Supervision, M.R.T.; Funding Acquisition, N.S. and M.R.T.

## Data and Code Availability

The accession number for the whole genome sequencing data reported in this paper for the EHE extended primary cell cultures is SRA: PRJNA1164735 The accession number for the bulk RNA-Seq data reported in this paper for the EHE extended primary cell cultures, EHE tumors, and human pulmonary artery endothelial cells is GEO: GSE277493 The accession number for the single cell RNA-Seq data report in this paper for EHE extended primary cell culture 001 is GEO: GSE277495

## Materials and Methods

### Primary Cell Culture Generation

EHE tumor samples were received from the EHE Biobank, an ethically-approved centralized biorepository. The protocol (number NB100040), informed consent, and supporting documents were reviewed and approved by North Star Review Board, a central Independent Review Board. All samples were de-identified and clinically annotated. Upon receipt, EHE tumor samples were cut into smaller pieces and treated overnight in a 37°C incubator with 1 mg/ml collagenase/dispase in DMEM media. The following day, media and tumor pieces were aspirated and placed in a 15mL conical tube for centrifugation. Media containing collagenase/dispase was discarded, and tumor pieces were placed back into the dish in complete endothelial cell growth media to allow cells to migrate out from the tumors over the course of 2 weeks, then subsequently passaged once confluent.

### Next-generation sequencing-based assay for detection of the *WWTR1*-*CAMTA1* gene fusion

Total RNA isolated from EHE 036 tumor was used for targeted library preparation using the Universal RNA Fusion Detection Kit for Ion Torrent (ArcherDX, Boulder, CO) and the Archer FusionPlex Sarcoma that uses anchored multiplex PCR as previously described ^60^. Following cDNA synthesis, the products were processed and ligated to adaptors. Two rounds of PCR were performed using a *CAMTA1* primer for target enrichment. The amplicon was clonally amplified and sequenced on the Ion Torrent personalized genomic machine (PGM) (ThermoFisher, Waltham, MA). Raw sequence data was analyzed with the Archer analysis pipeline version 3.3. Default thresholds and a minimum of five reads with three or more unique sequencing start sites to cross the breakpoint were set as the cutoff requirement to call the *WWTR1*-*CAMTA1* gene fusion.

### Cell culture

After collagenase/dispase treatment, EHE extended primary cell cultures were grown in EGM-2 Endothelial Cell Growth Medium-2 (EGM-2) BulletKit (catalog #: CC-3162) Lonza (Walkersville, MD, USA). EGM-2 was prepared with the provided amphotericin/gentacin omitted and was supplemented with 5% penicillin/streptomycin and to 15% fetal bovine serum (catalog #: A5670401) Gibco (Waltham, MA, USA). The EHE extended primary cell cultures were passaged 1/4 (25%) as cells reached 90% confluence. All cells were cultured at 37° C and 5% CO2. Human primary pulmonary artery endothelial cells (HPAEC), were obtained (PCS-100-022) from the American Type Culture Collection (ATCC, Manassas, VA, USA) and grown in EGM-2 with 15% fetal bovine serum and substituting 5% penicillin/streptomycin for the provided amphotericin/gentacin. Cells were passaged with 0.05% trypsin-EDTA.

### Compounds

VT-107 was obtained from MedChemExpress (Cat. No.: HY-134957). IK-930 was obtained from MedChemExpress (catalog #: HY-153585).

### Antibodies

Anti-TAZ antibody (catalog# HPA007415) utilized for western blot (1:2000) was obtained from Sigma-Aldrich (St. Louis, MO, USA). Anti-RNA Pol II antibody (4H8; catalog# 39097) (1:2000) was obtained from Active Motif. Anti-p16 antibody (EPR1473; catalog# ab108349) utilized for western blot (1:2000 in 5% BSA) was obtained from Abcam (Cambridge, MA, USA). Anti-β-actin antibody (AC-15; catalog #A544) utilized for western blot (1:10,000) was obtained from Sigma-Aldrich (St. Louis, MO, USA).

### Immunohistochemistry

CAMTA1 immunohistochemistry was performed on 4-µm-thick tissue sections after PT Link (Agilent Dako; Santa Clara, CA) deparaffinization, rehydration, and heat-induced epitope retrieval in HpH Target Retrieval Solution (Agilent Dako; pH 9) on an Autostainer Link 48 (Agilent Dako) using a rabbit polyclonal antibody raised against a recombinant protein corresponding to amino acids 451 to 535 of CAMTA1 (Novus Biologicals; Centennial, CO; product code NBP1-93620; 1:500 dilution; 30 minute incubation) and the polymer-based EnVision FLEX detection system (Agilent Dako; 30 minute incubation). An epithelioid hemangioendothelioma from the UIHC surgical pathology archive served as the positive tissue control, while a multi-tissue block including normal tonsil, placenta, lung, colon, colon cancer, and myometrial tissue served as the negative tissue control.

### qRT-PCR

EHE 001 and EHE 036 cells were scraped and collected using TRIzol reagent (Ambion-Life Technologies). Total RNA was isolated using the PureLink RNA Mini Kit (Invitrogen-ThermoFisher Scientific). On-column DNase (Invitrogen) treatment was performed according to manufacturer’s instructions. Purified RNA was quantified using a NanoDrop (ThermoFisher) and 1 µg was converted to cDNA using SuperScript III First-Strand Synthesis System (Invitrogen) according to manufacturer’s instructions. PCR amplification was performed in technical triplicates on the Applied Biosystems QuantStudio 3 Real-Time PCR System (Applied Biosystems-ThermoFisher). TaqMan Universal PCR Master Mix (Applied Biosystems-ThermoFisher) and PrimeTime standard qPCR primer/probe sets from Integrated DNA Technologies (Iowa City, IA, USA) were utilized. The qPCR cycling conditions were as follows: 95 °C10:00(95 °C0:15, 60 °C1:00)40. Relative quantitation was performed utilizing the delta-delta CT method and β-Actin CT values as the reference control. Each experiment was repeated at least twice. The following primers and probes were used:

EHE001-F-5’ GCTGGGAGATGACCTTCAC-3’ EHE001-R-5’ TGGAGAGGACCAGCTCTT-3’

EHE001-Probe 5’ AGGTACTTCCTCAACTCGGAGAGCC-3’

EHE036-F-5’ GCTGGGAGATGACCTTCAC-3’ EHE036-R-5’ TCCATCTGCTCCAGGGT-3’

EHE036-Probe 5’ ACTTCCTCAACCTCCTGCCGTC-3’

ActB-F-5’ ACCTTCTACAATGAGCTGCG-3’ ActB-R-5’CCTGGATAGCAACGTACATGG-3’ ActB-Probe 5’ ATCTTCTTCTCGCGGTTG-3’

### Poly-HEMA

A poly(2-<u>h</u>ydroxy<u>e</u>thyl <u>m</u>eth<u>a</u>crylate), or poly-HEMA, solution was made at 20 mg/mL in 95% ethanol. 96-well tissue culture plates were coated with 130 µL of poly-HEMA solution and UV sterilized overnight. Cells were plated at 4 × 10^3^ cells/100 µL per well and incubated at 37°C and 5% CO_2_. Proliferation was assessed every other day by adding 10 µL of Cell Counting Kit-8 reagent (Dojindo Molecular Technologies, Inc, Rockville, MD, USA). The plates were then incubated for 1, 2, and 4 hours at 37°C and 5% CO_2_ and then absorbance at 450 nm was measured using a Synergy H1 Hybrid Multi- Mode Microplate Reader (Biotek, Winooski, VT, USA).

### Proliferation Assay

Cells were plated at 1.5 × 10^3^ cells/100 µL per well and incubated at 37°C and 5% CO2. Proliferation was assessed every other day by adding 10 µL of Cell Counting Kit-8 reagent (Dojindo Molecular Technologies, Inc, Rockville, MD, USA). The plates were then incubated for 1, 2, and 4 hours at 37°C and 5% CO_2_ and then absorbance at 450 nm was measured using a Synergy H1 Hybrid Multi-Mode Microplate Reader (Biotek, Winooski, VT, USA).

### Western Blot

Cell pellets were lysed in radioimmunoprecipitation assay (RIPA) buffer, containing cOmplete Protease Inhibitor Cocktail (EDTA-free) (Roche) and PhosSTOP Phosphatase Inhibitor Cocktail (Roche) according to the manufacturer’s instructions. Total protein concentration was measured using Pierce BCA Protein Assay Kit (ThermoFisher Scientific, Waltham, MA, USA). Between 50 and 100 µg of total protein was loaded onto a gradient (4–15%) polyacrylamide gel (BioRad, Hercules, CA, USA). Proteins were then transferred to a polyvinylidene difluoride (PVDF) membrane and probed with antibodies described above.

### Nuclear and cytoplasmic fractionation

Nuclear and cytoplasmic fractionation was performed with the Nuclear Extract Kit (catalogue # 40010) from Active Motif (Carlsbad, CA). Cells collected from 15 cm cell culture plates were centrifuged and resuspended in 500 µL 1X Hypotonic buffer. After incubating for 15 min on ice, 25 µL of detergent was added to resuspended cells which were subsequently vortexed for 10 seconds. After centrifugation, the supernatant cytoplasmic fraction was removed and stored. The remaining nuclear pellet was resuspended in 50 µL complete lysis buffer followed by homogenizing with pestle for 5-10 seconds, briefly vortexing, and incubating on ice for 10 minutes. Homogenization, vortexing, and incubation on ice was completed an additional two times for a total of 3 times. After centrifugation, the supernatant nuclear fraction was removed and stored.

### Soft Agar

The base layer of 0.5% agarose was plated into 6-well plates (2 mLs/well) and allowed to solidify for 1 hour. 1 × 10^4^ cells/2 mL in 0.35% agarose was added to form the top layer (2mLs/well). Plates were left in a laminar flow hood and allowed to solidify at room temperature for 3 hours. Starting the following day, 1 mL of media was added to each well weekly for the duration of the experiment. Colonies were allowed to grow for 3-5 weeks at 37°C and 5% CO_2_ before imaging. Each experiment was repeated at least twice.

### Whole Genome Sequencing

Genomic DNA was isolated from the EHE 001, EHE 036, EHE 053, and EHE 061 extended primary cell cultures using the DNeasy Blood and Tissue Kits for DNA Isolation (Qiagen, Germantown, MD). Single samples from each extended primary cell culture were sequenced using an Illumina NovaSeq 6000 in the Iowa Institute of Human Genetics (IIHG) Genomics Core Facility as described above. Whole genome variant analysis was performed on the Dragen Bio-IT platform (Illumina, version SW: 07.021.645.4.1.5, HW: 07.021.645). The following flags were set: --intermediate-results-dir /staging --soft-read-trimmers polyg,adapter,n --trim-adapter-read1 /Shared/Bioinformatics/data/mchiment/adapters.fasta --trim-adapter- read2 /Shared/Bioinformatics/data/mchiment/adapters.fasta --enable-bam-indexing true --enable-duplicate-marking true --enable-map-align-output true --enable-map-align true --enable-variant-caller true.

### RNA Sequencing

Total RNA was isolated from biological triplicates of the EHE 001, EHE 036, EHE 053, and EHE 061 extended primary cell cultures using the PureLink RNA Mini Kit (Invitrogen-ThermoFisher Scientific). On-column DNase (Invitrogen) treatment was performed according to manufacturer’s instructions.

Transcription profiling using RNA-Seq was performed by the University of Iowa Genomics Division using manufacturer recommended protocols. Briefly, 100 ng of DNase I-treated total RNA was used to prepare sequencing libraries using the Illumina stranded total RNA prep, ligation with ribo-zero plus library kit (Cat. #20040529, Illumina, Inc., San Diego, CA). The molar concentrations of the resulting indexed libraries were measured using the Model 2100 Agilent Bioanalyzer (Agilent Technologies, Santa Clara, CA) and combined equally into a pool for sequencing. The concentration of the library pool was measured using the Illumina Library Quantification Kit (KAPA Biosystems, Wilmington, MA) and sequenced on the Illumina NovaSeq 6000 genome sequencer using 100 bp paired-end SBS chemistry.

Barcoded samples were pooled and sequenced using an Illumina NovaSeq 6000 in the Iowa Institute of Human Genetics (IIHG) Genomics Core Facility. Paired-end reads were demultiplexed and converted from the native Illumina BCL format to fastq format using an in-house python wrapper to Illumina’s ‘bcl2fastq’ conversion utility. FASTQ data were processed with nf-core/rnaseq (v3.12), a best-practices pipeline available at the open-source ‘nf-core’ project (https://nf-co.re/, Nextflow version 22.04) ^61^. Reads from the samples were aligned against the Ensembl reference ‘GRCh38’ using the STAR aligner.

Concurrently, reads were also pseudo aligned to the transcriptome using the Salmon aligner ^62^. Salmon performs its own internal quantitation, yielding estimated counts and values in length-normalized TPM (transcripts per million). Transcript-level abundances were converted to gene-level counts. Quality control was performed with MultiQC, a computational tool collates QC metrics to detect common QC problems. Gene-level counts were used for differential gene expression analysis with DESeq2 ^63^.

Bioconductor package ‘PCAExplorer’ was used for exploratory analysis ^64^. Venn diagrams were generated with the ‘gplots’ library from Bioconductor. Furthermore, the DE gene lists were analyzed using Advaita Bio’s iPathwayGuide (https://www.advaitabio.com/ipathwayguide) to detect and predict significantly impacted pathways. This software analysis tool implements the ‘Impact Analysis’ approach that takes into consideration the direction and type of all signals on a pathway, the position, role and type of every gene, ^65,66^. To assess overlap between data sets, hypergeometric analyses and Matthews correlation coefficient (MCC) calculations were carried out in R, version 4.4.1. The upper tail of the hypergeometric distribution was calculated using the phyper() function.

### Single Cell RNA-Sequencing

Samples from the EHE 001 extended primary cell culture contain 10,0000 cells from an early passage (p6) and late passage (p17) were submitted in biological duplicates. Barcoded samples were pooled and sequenced using an Illumina NovaSeq 6000 in the Iowa Institute of Human Genetics (IIHG) Genomics Core Facility. Paired-end reads were demultiplexed and converted from the native Illumina BCL format to fastq format using an in-house python wrapper to Illumina’s ‘bcl2fastq’ conversion utility. FASTQ data were pre-processed with the CellRanger (v7.1, 10X Genomics) pipeline using the “refdata-gex- GRCh38-2020-A” reference genome. Filtered barcode matrices were loaded into an R Studio environment and further analyzed with Seurat (v5). No downstream filtering for % mitochondrial expression or UMI and feature counts was performed owing to our desire to assess the cell lines in their entirety. Samples were integrated using the ‘RPCA’ method in Seurat. Clusters were detected with Seurat’s “FindNeighbors” (dims=1:30) and “FindClusters” (resolution=0.1) functions, and the UMAP projection was calculated for the integrated object. Loupe Browser (v7, 10X Genomics) was used for inspection of the integrated object.

### β-galactosidase staining

Early and late passage EHE 001 cells were plated in 6-well plates and stained the next day stained using the Senescence β-Galactosidase Staining Kit (Cell signaling Technology #9860, Danvers, MA, USA. Brightfield and phase contrast images were taken on Leica DM IL fluorescence and phase contrast microscope.

### Statistics

For proliferation, poly-HEMA, soft agar assays, statistical significance was evaluated student’s unpaired two-tailed *t*-test. To assess enrichment on RNA-Seq data sets, hypergeometric analyses and Matthews correlation coefficient (MCC) calculations were carried out in R, version 4.4.1. The upper tail of the hypergeometric distribution was calculated using the phyper() function. Each experiment was repeated at least twice. Error bars were used to define one standard deviation. For all panels, ****p<0.0001, *** p<0.001, ** p<0.01, *p<0.05.

---

**Figure S1.**
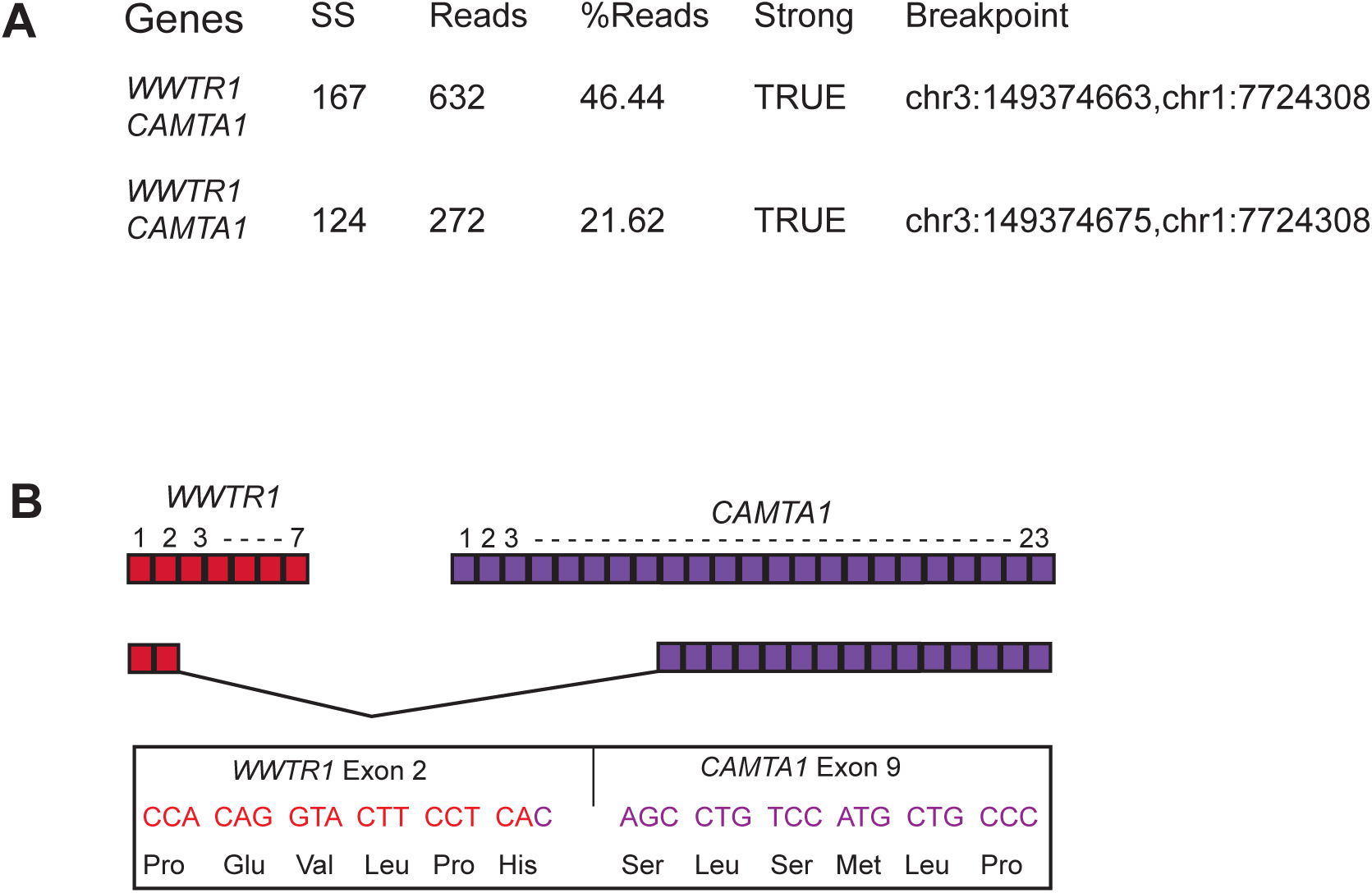
Schematic for RNA breakpoint for EHE 036. **(A)** Anchored multiplex PCR showing the relative frequency of the two *WWTR1*-*CAMTA1* fusion transcripts identified in EHE 036. (**B**) Diagram showing the breakpoint for the less abundant fusion transcript in (**A**).

**Figure S2.**
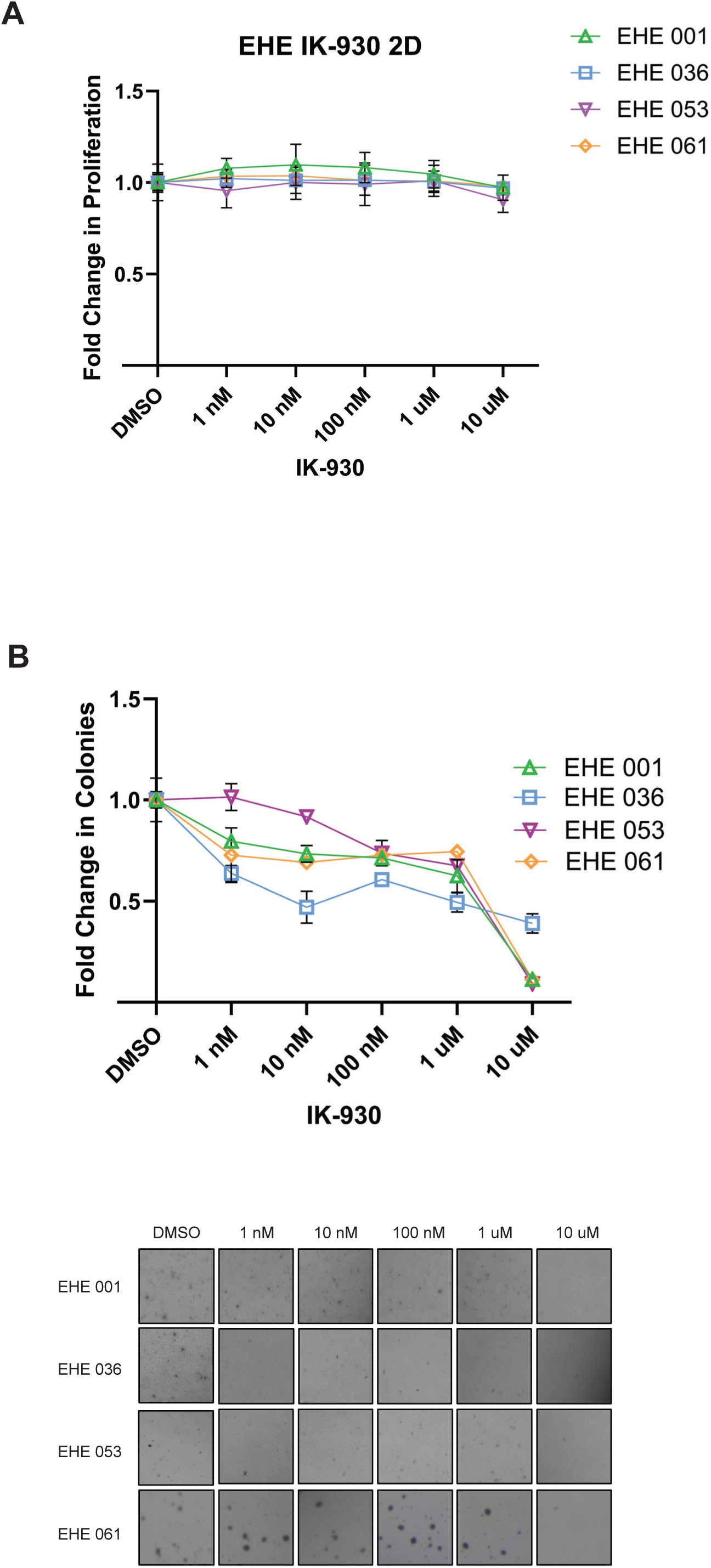
Effect of IK-930 on EHE extended primary cell cultures. **(A)** IK-930 does not significantly affect proliferation in EHE cell cultures. (**B**) IK-930 decreases anchorage independent growth in EHE cell cultures.

**Figure S3.**
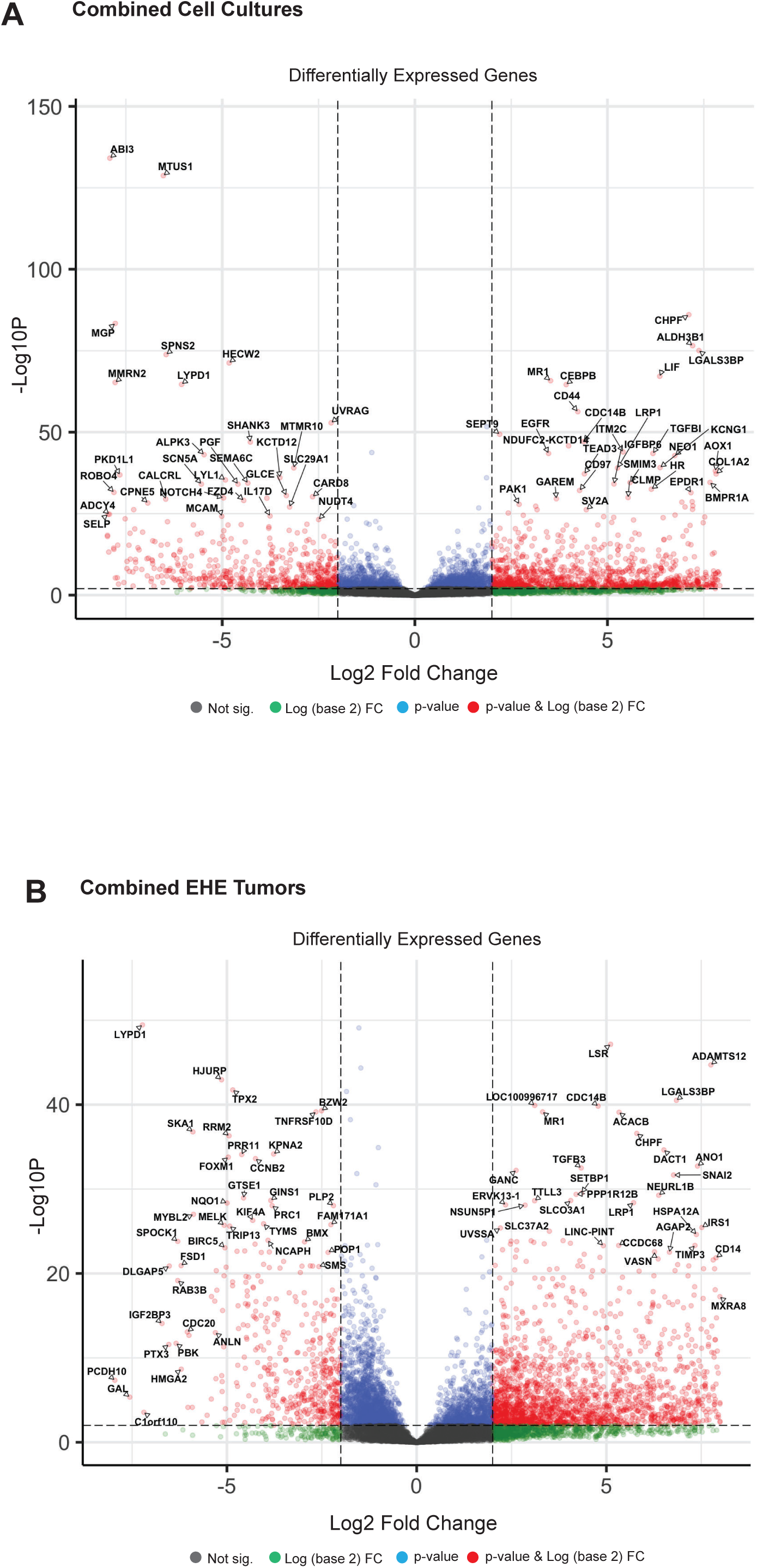
Differentially expressed genes for EHE extended primary cell cultures and EHE tumors. Volcano plots of differentially expressed genes for **(A)** the combined EHE cell cultures and **(B)** Combined EHE donor tumors. Differentially expressed genes normalized to HPAEC gene expression.

**Figure S4.**
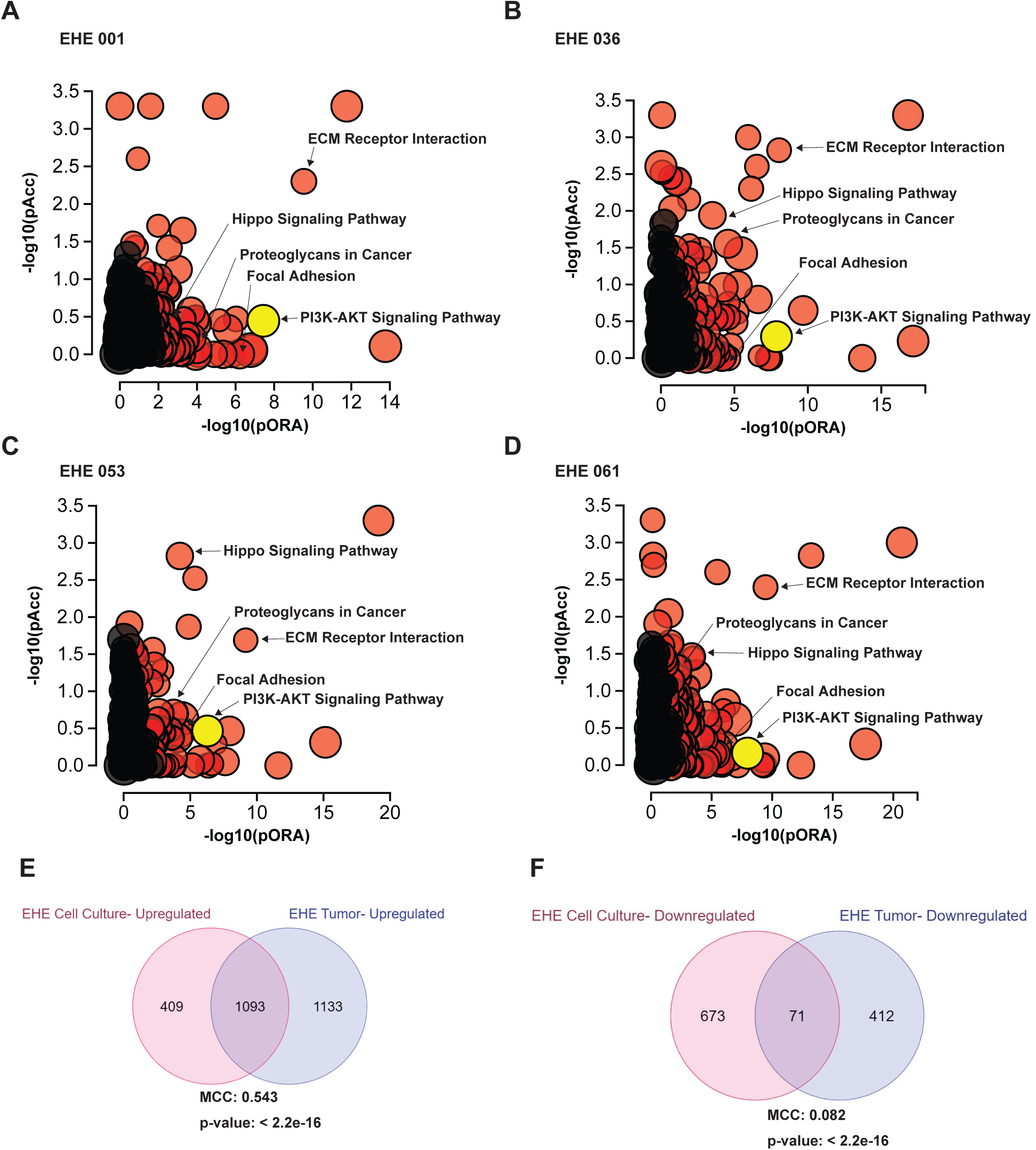
Gene expression in individual EHE extended primary cell cultures. Bulk total RNA-Seq is performed on the EHE 001, EHE 036, EHE053, EHE061 and HPAEC cell cultures; gene expression in EHE extended primary cell cultures is normalized to HPAEC gene expression. (**A-D**) Scatter plots of pathway enrichment analysis showing probability of over-representation (pORA) and probability of accumulation (pAcc) as calculated by iPathwayGuide; -log10(pAcc) is graphed vs -log10(pORA); for EHE 001 (**A**), for EHE 036 (**B**), for EHE 053 (**C**), and for EHE 061 (**D**). (**E**) Venn diagram showing overlap of upregulated genes in EHE extended primary cell cultures (combined) vs. upregulated genes in EHE tumors (combined). (**F**) Venn diagram showing overlap of downregulated genes in EHE extended primary cell cultures (combined) vs. downregulated genes in EHE tumors (combined). To assess enrichment, hypergeometric analyses and Matthews correlation coefficient (MCC) calculations were carried out in R, version 4.4.1. The upper tail of the hypergeometric distribution was calculated using the phyper() function.

**Figure S5.**
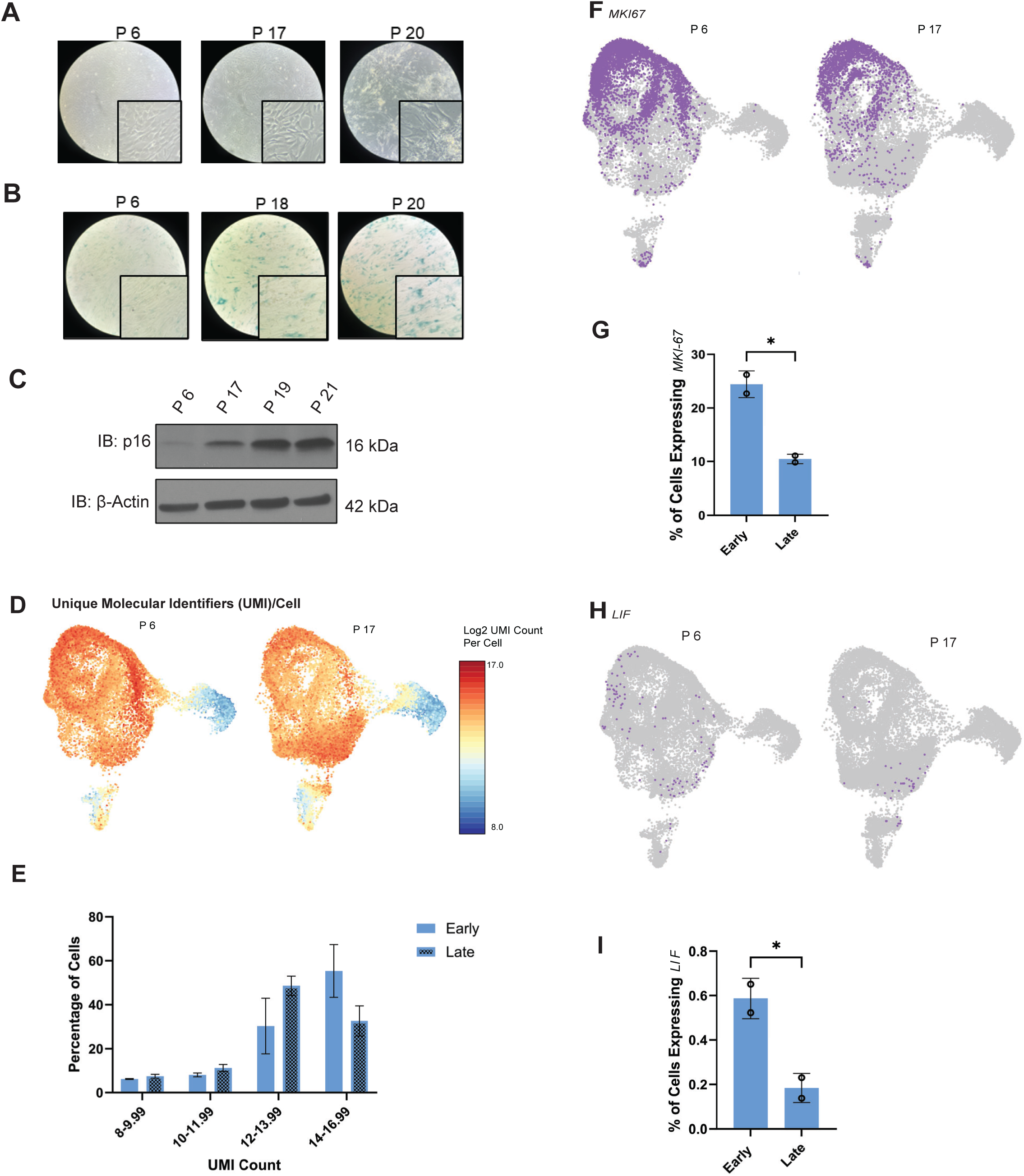
Defining different subsets within the EHE extended primary cell cultures. (A) Brightfield image showing changes in cell morphology in EHE 001 as a function of passage number. (B) β-Galactosidase staining in early and late passage EHE extended primary cell cultures. (**C**) p16 expression with continued passage of EHE 001 by western blot. (**D**) Unique molecular identifiers (UMI)/cell in early passage (p6) vs. late passage (p17) EHE 001 cells. (**E**) Graphical representation of unique molecular identifiers (UMI) in early passage (p6) vs. late passage (p17) EHE 001 cells. Percentage of cell graphed as a function of Log2UMI count per cell. (**F**) *MKI67* expression in early (p6) vs. late passage (p17) EHE 001 cells. (**G**) Graphical representation of MKI67 expression in early (p6) vs. late passage (p17) EHE 001 cells. (**H**) *LIF* expression in early (p6) vs. late passage (p17) EHE 001 cells. (**I**) Graphical representation of *LIF* expression in early (p6) vs. late passage (p17) EHE 001 cells. For expression of *MKI67* and *LIF*, statistical significance was evaluated with unpaired two-tailed Student’s *t*- test. Error bars were used to define one standard deviation. For all panels, ****p<0.0001, ***p<0.001, **p<0.01, *p<0.05.

